# ScAN1.0: A Reproducible and Standardized Pipeline for Processing 10X Single Cell Rnaseq Data

**DOI:** 10.1101/2022.11.07.515546

**Authors:** Maxime Lepetit, Mirela Diana Ilie, Marie Chanal, Gerald Raverot, Philippe Bertolino, Christophe Arpin, Franck Picard, Olivier Gandrillon

## Abstract

Single cell transcriptomics has recently seen a surge in popularity, leading to the need for data analysis pipelines that are reproducible, modular, and interoperable across different systems and institutions.

To meet this demand, we introduce `scAN1.0`, a processing pipeline for analyzing 10X single cell RNA sequencing data. `scAN1.0` is built using the Nextflow DSL2 and can be run on most computational systems. The modular design of Nextflow pipelines enables easy integration and evaluation of different blocks for specific analysis steps.

We demonstrate the usefulness of `scAN1.0` by showing its ability to examine the impact of the mapping step during the analysis of two datasets: (i) a 10X scRNAseq of a human pituitary gonadotroph tumor dataset and (ii) a murine 10X scRNAseq acquired on CD8 T cells during an immune response.

## 2 Introduction

The recent surge in single cell transcriptomics, particularly through singlecell RNAseq (scRNAseq), has led to the need for data analysis pipelines that are reproducible, modular, and able to interoperate across different systems and institutions.

The first step in scRNAseq data analysis consists in generating a count matrix from `FASTQ` sequence files. However, this step is often overlooked although it can be critical for the success of the analysis. Therefore, it is important to have access to analysis pipelines that allow for easy verification of the impact of various analysis steps, such as the nature of the Gene Transfer Format (`GTF`) file used or the normalization method on the generation of the count matrix.

scRNAseq analysis packages such as Seurat [1] are designed to be userfriendly, but do not offer easy options for incorporating alternative low-level analysis steps. To address this challenge, we developed `scAN1.0`, a processing pipeline for 10X scRNAseq data that can be run on most computational systems using the Nextflow DSL2.

Nexflow [2] is a bioinformatics workflow management tool that aims to improve reproducibility, modular architecture, and cross-environment compatibility in bioinformatics pipelines, through:

1. Reproducibility: Nexflow allows users to easily and accurately reproduce their pipelines by providing a clear and traceable record of all the steps that were taken and the input and output data for each step. This helps to ensure that the results of a pipeline can be easily reproduced and validated by other researchers.
2. Modular architecture: Nexflow has a modular architecture, which means that pipelines can be built by combining smaller, reusable components called “tasks”. This allows users to easily reuse tasks in multiple pipelines and makes it easier to maintain and update pipelines over time.
3. Cross-environment compatibility: Nexflow is designed to be compatible with a wide range of computing environments, including on-premises clusters, cloud environments, and desktop computers. This makes it easier for users to run pipelines in the environment that best suits their needs and enables them to easily scale their pipelines as their needs change.

We illustrate the benefit of using `scAN1.0` by showing its ability to assess the impact of the mapping step on the resulting UMAP projection as well as on some specific gene identification using two datasets: (i) a 10X scRNAseq of a human pituitary gonadotroph tumor dataset and (ii) a murine 10X scRNAseq acquired on CD8 T cells during an immune response.

## 3 Results

### 3.1 Version annotation effect

We first assessed the impact of the Ensembl version of the `GTF` file on the final output using the human Gonadotroph tumour dataset as an input (see Material and methods). `GTF` files and their corresponding `FASTA` files were downloaded via `ftp` protocol using the --version parameter (see section 5). We set Cellranger as the default mapper and then assessed the impact of 4 different annotation releases (93, 98, 103 and 106) on the number of detected genes (figure 2A) and on the count per genes as assessed with the CHGA gene (Figure 2B).

**Figure 1:**
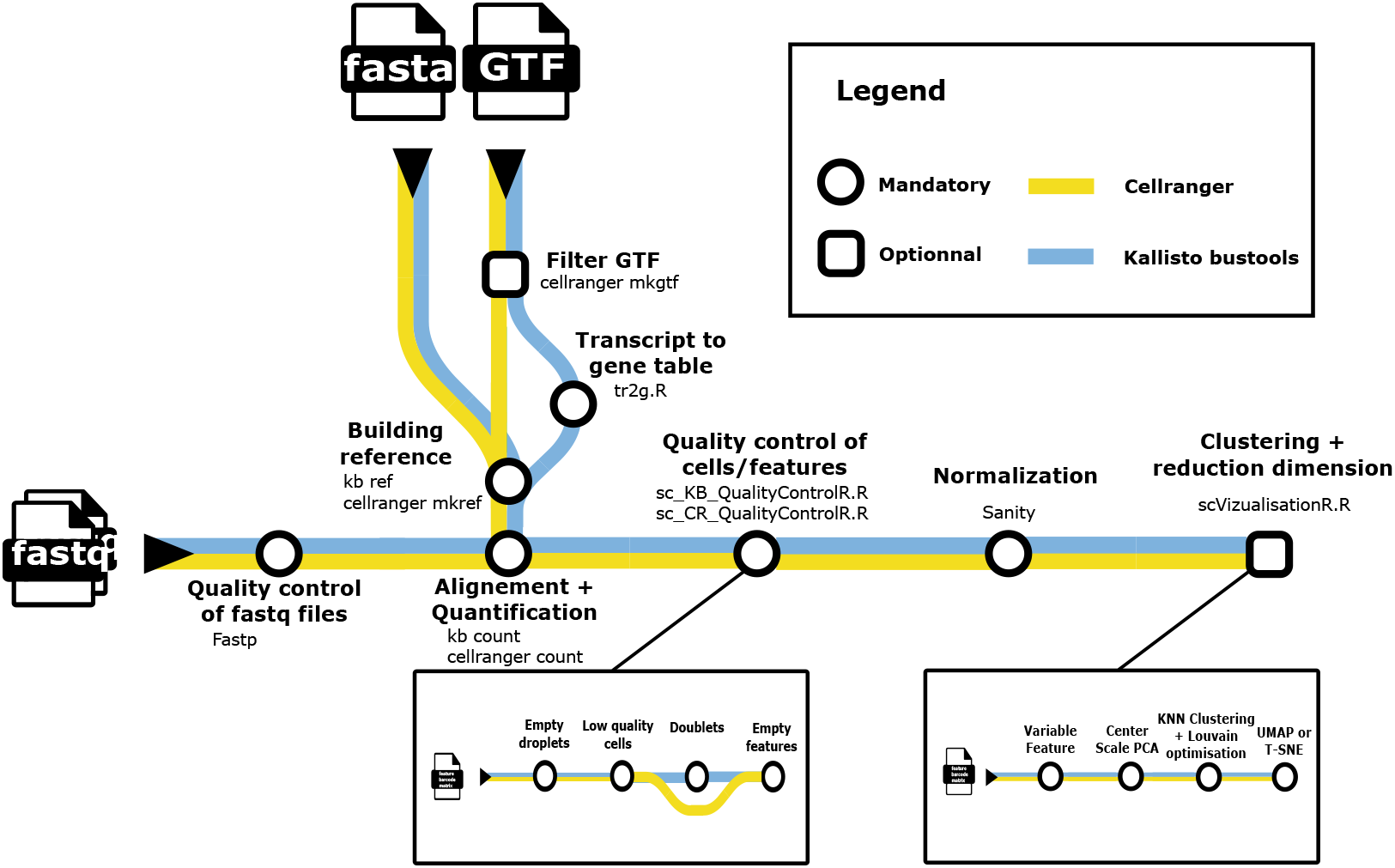
A metromap view of the `scAN1.0` pipeline. Tools used for the processing steps are given below the name of the process.

**Figure 2:**
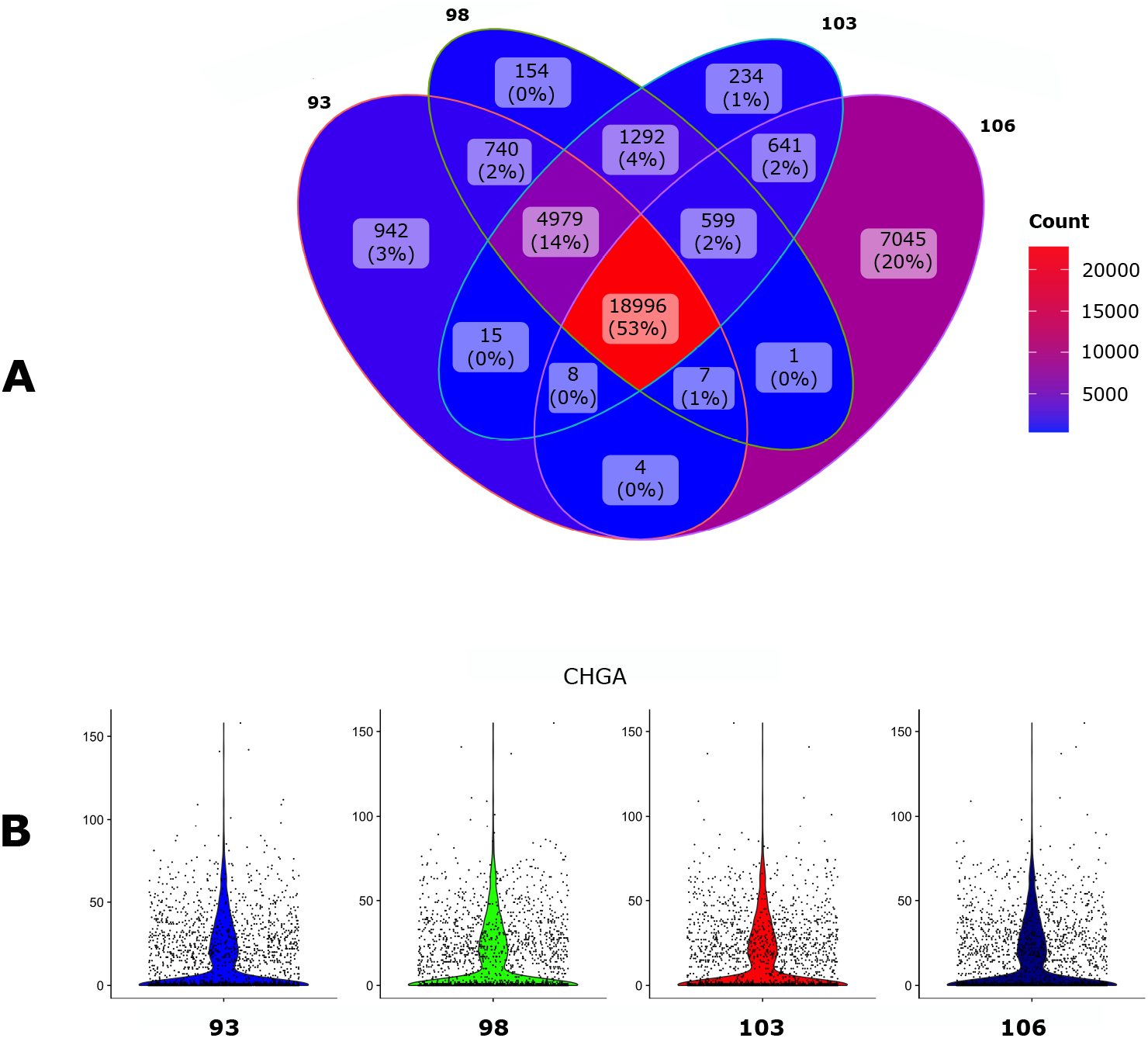
Impact of the `GTF` version. The human gonadotroph tumors dataset was submitted to `scAN1.0` with 4 different versions of the `GTF` for mapping with Cellranger. A. Venn diagram of the number of the detected genes when using the different versions. B. Violin plot representation of the impact of the `GTF` version on the UMI counts for the CHGA gene.

The overall impact of the `GTF` version seems relatively modest, especially in regard to the number of UMI counts for CHGA. However, the 106 version allowed to identify a larger number of genes and was kept for the next step.

### 3.2 Comparing filtered with unfiltered annotations

We then assessed the impact of filtrating the `GTF` file with the `cellranger mkgtf` function. This filtration step is intended to remove unwanted genes classified by biotype, such as long non-coding RNAs (https://www.ensembl.org/info/genome/genebuild/biotypes.html). We used the default values of that function that removes biotypes such as gene_biotype:pseudogenes from the `GTF` annotation file.

This step reduces the presence of multi-mapping due to non-regularly processed nucleic acids, removing ambiguity about gene identification and allowing quantification of properly mapped sequences.

We observed that this filtration step indeed had a major impact on both the number of genes detected (Figure 3A) but also on gene counts (Figure 3B).

**Figure 3:**
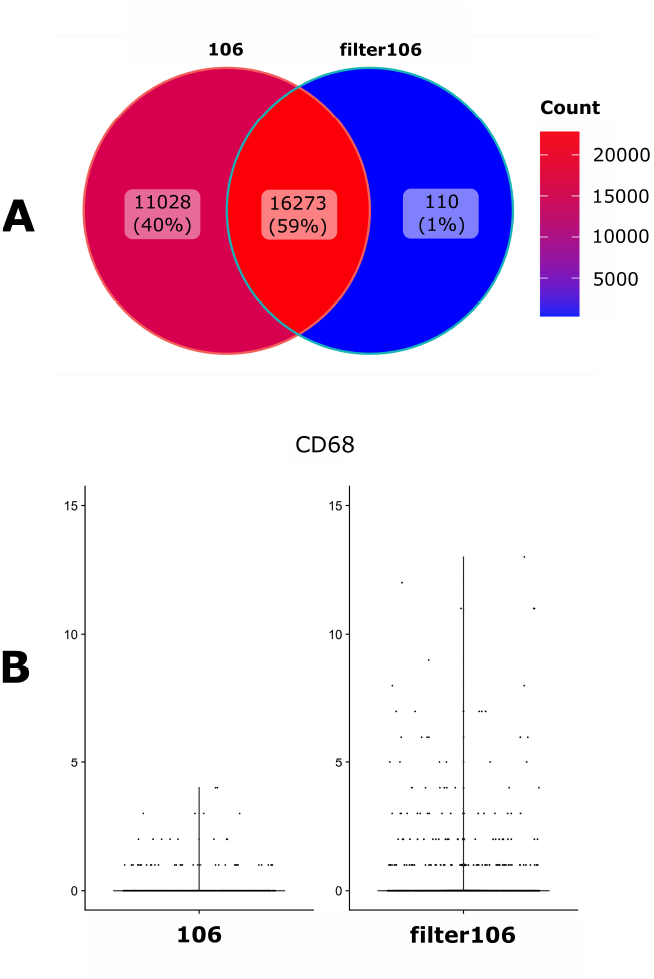
Impact of the `GTF` filtration. A. Venn diagramm of the number of the detected genes when using either an unfiltered (106) of filtered (filter106) version of the `GTF` file. B. Violin plot representation of the filtration impact on the UMI counts for the CD68 gene.

### 3.3 Cellranger versus Kallisto-bustools

We then compared the impact of two popular alignment tools for single-cell RNA sequencing (Kallisto-bustools and Cellranger) using the 10x Genomics pre-built Cellranger reference packages version 2020-A for human.

As seen in Figure 4A, 81 % of the genes were identified by both algorithms whereas Kallisto-bustools identified more genes than Cellranger.

**Figure 4:**
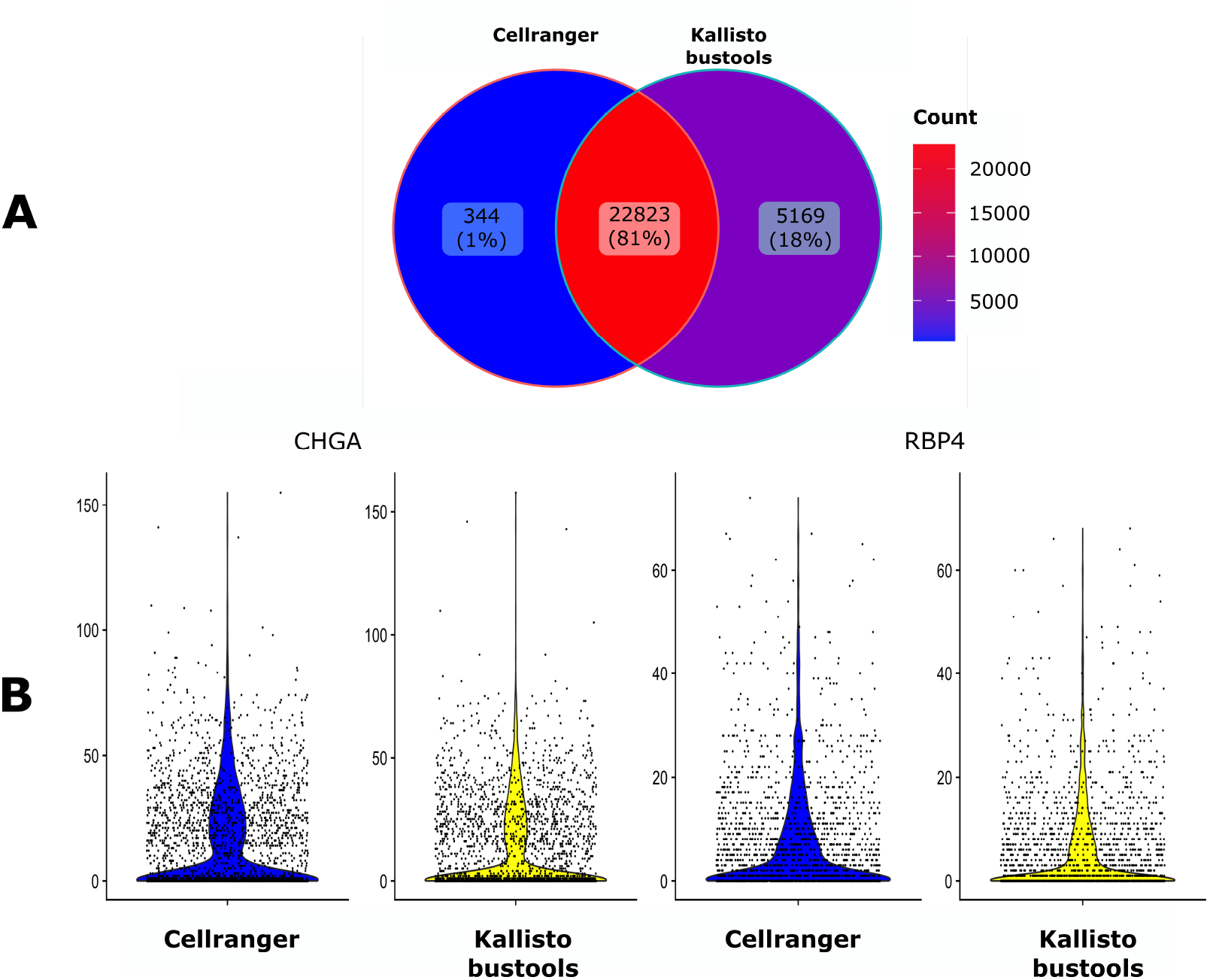
Impact of the mapping tool. A. Venn diagram of the number of the detected genes when using either Cellranger (CR) or Kallisto-bustools (KB) as an alignment tool. B. Violin plot representation on the UMI counts for two genes, CHGA and RBP4.

The impact of the mapper on counts for specific genes seemed to be negligible (Figure 4B). Therefore this tends to favor Kallisto-bustools for downstream analyses.

We also assessed the impact of the mapper choice on the final clustering step. As seen in Figure 5, the impact was modest but apparent (e.g. cluster number 1 in the Cellranger dataset was split in two in the Kallisto-Bustools dataset).

**Figure 5:**
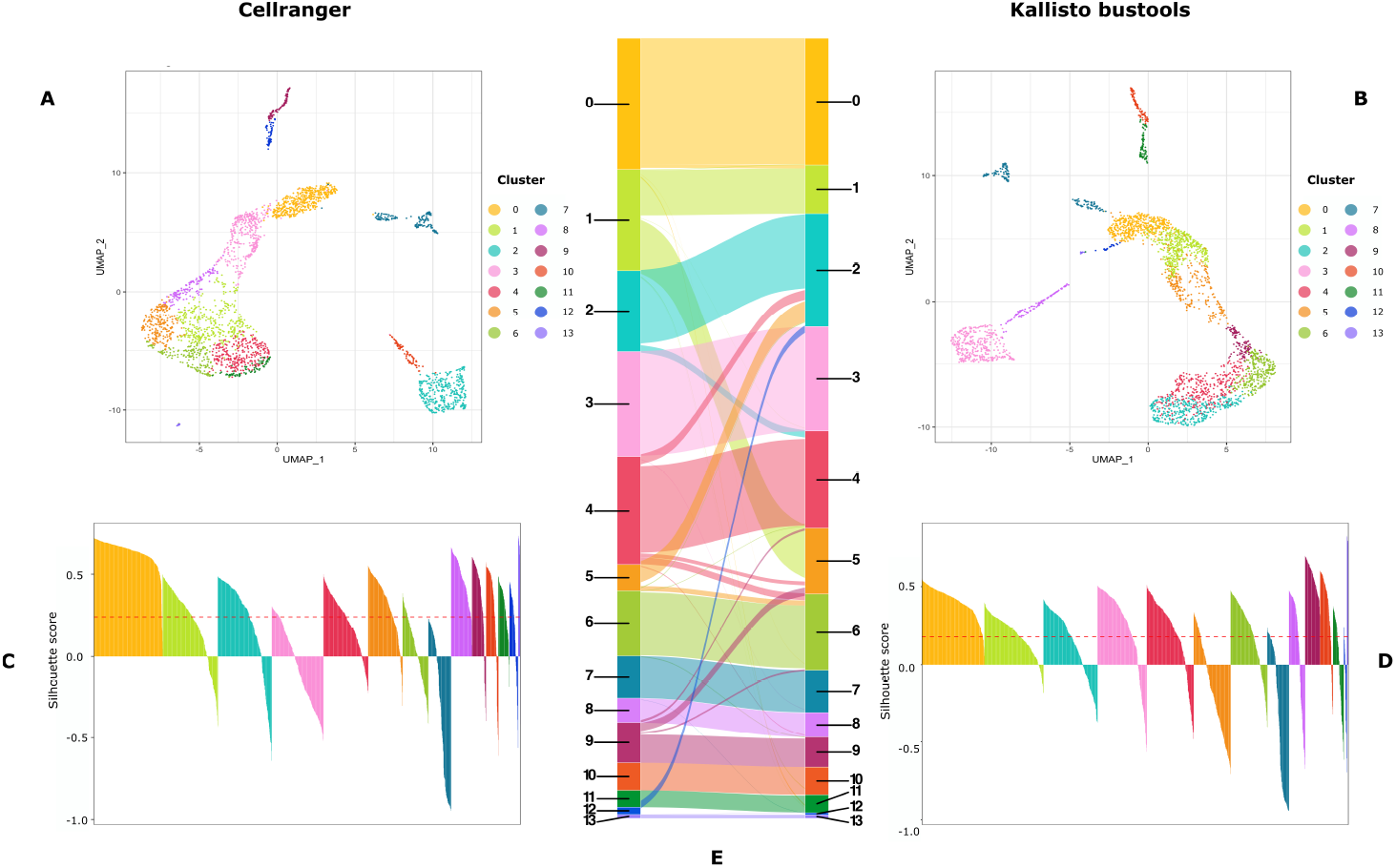
Cluster characterization: UMAP representation (A and B) and Silhouette scores (C and D) of the clusters obtained on data processed with CellRanger (A and C) or Kallisto-bustools (B and D). In E, is shown an alluvial plot highlighting the conservation and differences in cluster composition depending upon the initial mapping method.

Please, note that the different positioning of the clusters on each UMAP is due to known flaws associated with a 2D representation using UMAP (see e.g. [3])

### 3.4 Application of scAN1.0 to a CD8+ T cell murine dataset

The final combination that was found to be the most effective for our human dataset therefore was using Kallisto-bustools together with the filtered version of the 106 `GTF`.

We decided to apply `scAN1.0` to a published scRNA-seq dataset acquired from murine individual CD8+ T cells throughout the course of their differentiation in response to viral infection [4] and in order to compare the two alignment tools Kallisto-bustools and Cellranger.

As seen in Figure 6, overall less genes were detected than observed in human cells with 11587 genes identified by both algorithms. Quite interestingly we observed the very same increase (plus 20 %) in in the number of genes identified by Kallisto-bustools when compared with Cellranger.

**Figure 6:**
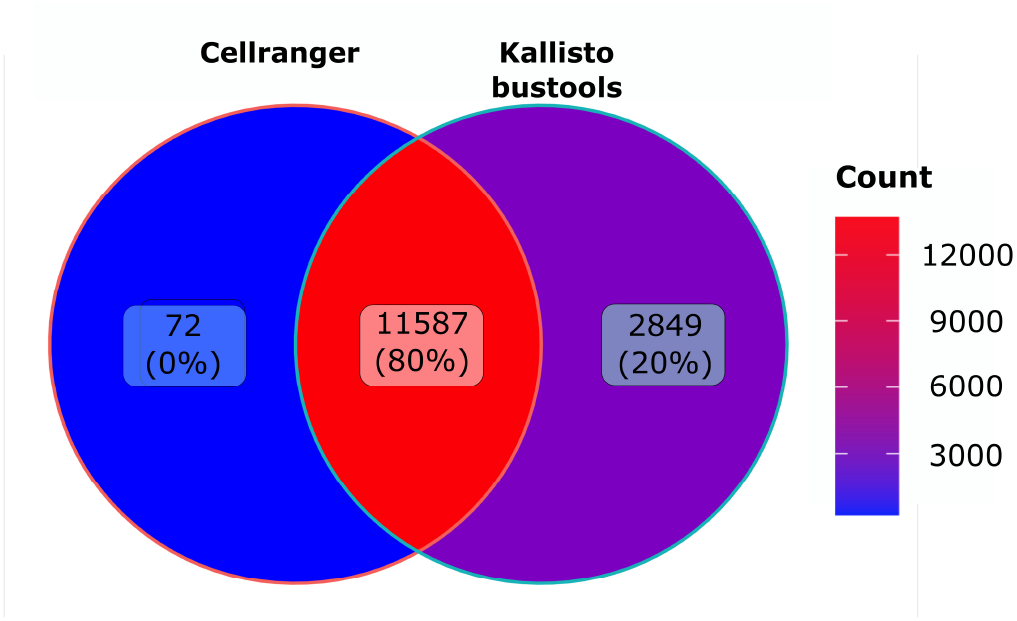
Murine dataset analysis: Venn diagramm of the number of genes detected using Cellranger (CR) or Kallisto-bustools (KB) as alignment tool the murine CD8 dataset.

## 4 Discussion

We described `scAN1.0`, a Nextflow based processing pipeline of 10X scR-NAseq data.

By applying `scAN1.0` to a tumor dataset, we showed that the impact of the annotation version was relatively modest although using the latest Ensembl release (106) of the `GTF` and `FASTA` allows to identify a larger number of genes.

As expected, filtrating the `GTF` files by removing unwanted genes based on 10X reference package of biotypes had a major impact both on the number of genes but also on gene counts. Furthermore, a recent study showed that `GTF` filtration led to a diminution of the number of processed pseudogenes and to a better identification of mitochondrial genes [5]. We therefore strongly recommend to use a filtered annotation when controlling data for mitochondrial content.

Futhermore, when using Kallisto-bustools instead of Cellranger the impact of the count numbers for specific genes seemed to be small but meaningful. Finally, Kallisto-bustools produced higher total number of genes detected than Cellranger on two very different datasets. For the human dataset, we obtained 5169 unique genes detected by Kallisto-bustool as compared to 344 unique genes detected by Cellranger. For the murine dataset, the ration was 2849 to 72.

From a biological standpoint, this represents a very significant increase in the amount of biological information at hand, and in our view, it fully justifies the use of Kallisto-bustools instead of Cellranger.

Thus, `scAN1.0` can be easily used to reproducibly compare different conditions of the pre-processing steps of scRNAseq analyses. Its modular composition of independent processes communicating via channels allows easy workflow pipeline modifications with DSL2 syntax and addition of new processing modules. For instance, we intend to include different normalization procedures in the future:

- The basic global-scaling normalization method from Seurat [6].
- `Sctransform` which uses regularized negative binomial regression and computes Pearson residuals that correspond to the normalized expression levels for each transcript [7].
- `Scran` which uses pooling-based size factor estimation [8].

The use of `scAN1.0` should be made straightforward to assess a combination of low-level steps together with the normalization step on the resulting output.

Up to now, two solutions were offered to anyone wanting to process scR-NAseq data: (i) using a turnkey solution like Seurat or (ii) chaining its own combination of required scripts (see e.g. [9]). Our solution bridges the gap between those two options: (i) it can be run easily in a ‘‘full Seurat mode” or (ii) it can used as it stands for exploring the impact of mapping solutions or (iii) it can be used for more advanced bioinformaticians to plug-in and easily chaining new algorithms.

One major limitation of using Nextflow is a lack of interactivity during pipeline running, which would allow the user to modify parameter on the fly. Instead, all pipeline parameters need to be defined and set in the launching command.

In conclusion, we believe that `scAN1.0` will be a useful tool for the growing community of 10X scRNAseq *aficionados*.

## 5 Material and methods

### 5.1 Pipeline description

Figure 1 describes the overall processing of the sequencing files with the ordering of all steps described in subsection 5.5.

### 5.2 Gonadotroph tumour dataset

#### Single cell preparation and sequencing

A tumor fragment from a gonadotroph surgically-resected adenoma was collected in Dulbecco’s Modified Eagle Medium (DMEM, cat 41965062; Life Technologies). Single-cell suspension of the resected fragment was obtained through mechanical and enzymatic dissociation (Collagenase P, cat 11213865001) followed by filtration through a 100 μm mesh-strainer (#732-2759, VWR international).

Red blood cells were eliminated using a 10-minute incubation with a commercial red blood cell lysis buffer (eBioscience, cat #00-4300-54). The whole process was achieved within the 2 hours following the surgical resection, cell viability was evaluated to reach at least 70 percent prior encapsulation. Generation of the library was done using a Chromium controller from 10xGenomics. The entire procedure was achieved as recommended by the manufacturer’s for the v3 reagent kit (Chromium Next GEM Single Cell 3’ Kit v3.1 (10xGenomics cat1000269). Single cell suspension was loaded onto a Chromium Single Cell A Chip, aiming for 5,000 cells in total. The cDNA was amplified after a reverse transcription step, followed by a cleaning using SPRIselect (Beckman Coulter), a quantification step, and enzymatic fragmentation before being sequenced using a NextSeq500 system (Illumina). `FASTQ` files were generated from sequencer’s base call files (BCLs) for each flowcell directory using `cellranger mkfastq`.

#### Study ethic approval

This work is part of the SPACE-PIT study (MR004 n21-5439). It was approved by the Hospices Civils de Lyon ethical committee and registered at the “Centre National Information et Liberté” (`CNIL.fr`) under the reference 20_098. Informed consent was obtained from the patients.

#### Processing

The dataset was processed using `scAN1.0` with the following parameters:

- species = human (default option)
- min_cells= 3
- min_feature_RNA = 500
- max_feature_RNA = 7000
- max_percent_mito= 25

The following options were used to choose either KB or CR:

- version =106
- quantif = cellranger or quantif = kb

The following options were used for filtrating `GTF`:

- filter`GTF`

The following options were used to choose `GTF` version:

- version = 106
- version = 103
- version = 98
- version = 93

The number of principal components of the PCA used for UMAP embedding and clustering was set to 10 as determined by the rule of thumb heuristic and the broken stick method [10]. The clustering resolution was set to 0.7.

### 5.3 CD8 T cells dataset

#### Single cells preparation and sequencing

We downloaded reads from GEO (GSE131847, https://www-ncbi-nlm-nih-gov.proxy.insermbiblio.inist.fr/geo/) with fastqdump (https://rnnh.github.io/bioinfo-notebook/docs/fastq-dump.html) using the --split-files option. Quality control of reads was performed with fastp (https://github.com/OpenGene/fastp) and resulting `FASTQ` files were submitted to `scAN1.0`.

To briefly describe this dataset (see [4] for the full description), P14 transgenic CD8 T cell, which recognise an LCMV epitope, were transferred to histo-compatible hosts that were acutely immunised the day after with the virus. Six days after infection, single responding P14 CD8 T cells from the spleens of immunised hosts were sorted and loaded into Single Cell A chips for partition into Gel Bead In-Emulsions in a Chromium Controller (10x Genomics). Single-cell RNA libraries were prepared according to the 10x Genomics Chromium Single Cell 3 Reagent Kits v2 User Guide and sequenced (paired-end) on a HiSeq 4000 (Illumina).

#### Processing

The dataset was processed using `scAN1.0` with the following parameters:

- version = 106
- species = mouse
- filter`GTF`
- min_feature_RNA = 500
- min_cells= 3
- max_percent_mito= 5
- chemistry V2

The following options were used to choose either KB or CR:

- quantif = cellranger or quantif = kb

### 5.4 Implementation

The `scAN1.0` pipeline was implemented using the reactive workflow manager Nextflow [2], coding with the DSL2 syntax extension. Nextflow simplifies the writing of computational pipelines by making them portable, scalable,parallelizable and ensuring a high level of reproducibility. Nextflow provides native support for container technologies such as Docker or Singularity. Each process in the pipeline will be run in a container. A reproducible container environment is built for each process from Docker images stored on the DockerHub. The analyses can be run on the user’s preferred computing platform. Using the configuration file and corresponding profile, the pipeline can be run on a local computer via Docker or Singularity, as well as on a high-performance computing (HPC) cluster or in cloud-based environments. In the case the workflow cannot complete all steps, a cache-based pipeline resume feature allows to recover the processed steps and the blocked process for debugging before resuming the workflow.

### 5.5 Input

`scAN1.0` takes three mandatory parameters as input: the paired-end `FASTQ` files from the 10X Chromium sequencing and two genomics files (one `FASTA` and one `GTF`).

`FASTQ` files store the nucleotide sequence and the associated sequencing quality scores. For paired-end sequencing, a `FASTQ` file is provided in a compressed `gzip` format for each read (R1 and R2). In the case a sample has been sequenced on several lanes, all R1 and R2 reads are concatenated together.

The mapping steps requires the input of two additional files corresponding to the species of interest:

- A `FASTA` genomic file which stores the raw genome sequence.
- A `GTF` file which stores genome annotation including gene positions.

One should note that for human datasets, `FASTA` and `GTF` files can be downloaded automatically by specifying a version number as an entry parameter (--version) available on the ENSEMBL database.

### 5.6 Preprocessing & Mapping

First, `FASTQ` files are processed and trimmed by using `Fastp` v0.20.1, an ultra-fast `FASTQ` preprocessor with useful quality control and data-filtering features [11]. Reads with phred quality (https://en.wikipedia.org/wiki/Phred_quality_score) at least equal to 30 (-q 30) are qualified for the quantification step. Length filtering is disabled (-L) while adapter sequence auto-detection is enabled (--detect_adapter_for_pe). To reduce overlapping annotation, we recommend and implemented (parameter --filter`GTF`) an optionnal tool of Cellranger (`mkGTF` function) that allows the `GTF` filtration based on biotype attributes of sequences. (see: https://support.10xgenomics.com/single-cell-gene-expression/software/release-notes/build and https://www.ensembl.org/info/genome/genebuild/biotypes.html)

The processed files are then mapped to a reference genome in order to quantify gene expression. In `scAN1.0`, the user can specify two different mappers (see below): Kallisto-bustools v0.26.0 [9] or Cellranger v5.0.1 [12].

- Kallisto-bustools is used thought the Python wrapper: kb_python. Starting with FASTA and `GTF` files, an index of the reference can be built as a colored De Bruij graph with Kallisto via kb ref. with default parameter. Once an index has been generated or downloaded, `kb count` uses Kallisto to pseudoalign reads and `bustools` to quantify the data.
- Cellranger creates and prepares a reference package with `cellranger mkref` function. The alignment is then run via `cellranger` count with default parameters as described on 10xgenomics.com.

### 5.7 Quality control

#### Empty Droplets

With droplet-based technologies, most of the barcodes in the matrix correspond to empty droplets (e.g., barcodes with the sum of expressions over all genes being null). They must be removed from Kallisto-bustools generated gene expression matrices. Thus, the Kallisto-bustools outputs were imported into `R` with a customized `R` function. The UMI total counts were ranked using `barcodeRanks()` function from `DropletUtils` v1.14.2. Empty droplets were removed by selecting the inflection point value on the resulting knee plot (lower cutoff = 10).

On its side, Cellranger handles empty droplets by itself and matrices cleared of empty droplets were directly imported from the standard filtered barcode output.

After importation, either gene expression matrices were converted as Seurat object (Seurat v4.0.4) with `CreateSeuratObject` function, including features detected in at least 3 cells (min.cells =3).

#### Low quality cells

On the resulting empty droplet-free matrices, the pipeline computes 3 QC metrics per sample: the number of unique features, the number of UMIs and the percent of mitochondrial gene counts; per cell. These QC metrics are then used to discard three main types of low quality cells [13]:

1. Cells in apoptosis may exhibit high % mitochondrial and low number of UMIs per cell.
2. Bad library preparation leads to low number of unique gene counts and low number of UMIs per cell.
3. Cellular doublets in droplets leads to high number of UMI and unique gene count.

For this, a threshold can be set by the user on `scAN1.0` parameters min_feature_RNA, min_ncount_RNA, max_percent_mito, max_feature_RNA, max_ncount_RNA; which default values are 500 and 0 for the first two and ‘‘adaptive” for the others. With the ‘adaptive” configuration, `scAN1.0` defines a certain number of median absolute deviations (MADs) away from the median to define maximum values *(median + 3*mad*) [14].

Furthermore, the pipeline uses the `R` package `DoubletFinder` v2.0.3 to detect and remove the potential doublets from the dataset. See [15] and [16].

#### Non-expressed genes

Genes with sum count along cells equal to 0 (e.g. not-expressed genes) are removed.

### 5.8 Normalization

After selection of high quality single-cells droplets and significantly expressed genes, the count matrices are then normalized using `Sanity`, a Bayesian algorithm to infer gene-expression state [17]. We are fully aware that the normalization of single cell transcriptomic data is a research field on its own (see e.g. [18] and citations therein), and we expect this block in the pipeline to be susceptible to be modified in future versions of the pipeline. The modularity of the Nextflow syntax makes it ideal for such additions.

### 5.9 Clustering and two-dimensional visualization

A final step consists in variable features selection with `Seurat::FindVariableFeatures` using the `vst` method (selection.method =“vst”) and selecting the 2000 first highly variable genes (nfeatures = 2000), followed by a first linear dimensionality reduction using PCA (Seurat::RunPCA with default parameter). The m first axis of the PCA are then used for the nearest-neighbor graph construction with `Seurat::FindNeighbors` function (dims=1:m).

Cluster determination was performed using the Louvain algorithm [19] run with `Seurat::FindClusters` function. The resolution parameter set by default to 0.7. The quality of the clustering was assessed using the Silhouette score [20]. Finally, non-linear dimensional representation (using either t-SNE [21] or UMAP [22]) is then performed using `Seurat::RunTSNE` or `Seurat::RunUMAP` with default parameters using the same number of dimensions than the nearest-neighbor graph building.

The resolution parameters used for clustering and the number of principal components kept for both clustering and dimension reduction embeddings can be respectively modified by the user at the start of the pipeline by the --resolution-clustering and the --principal-component parameters (default values: 0.7 and 10, respectively).

Alternatively, this last step can be skipped allowing the user to use their own clustering method. Similarly, to avoid the introduction of layers of complexity and simplify the pipeline usage, the automatic annotation of clusters was not introduced. Users can annotate their dataset manually.

### 5.10 Output

The complete list of process outputs is available on the Readme file of the gitlab repository (see section 6).

By default, all outputs are saved in the results directory, which is structured into subdirectories that are named according to each process. Results and graphics are stored in the subdirectories that correspond to each process. For the post-quantification steps, `scAN1.0` provides a Seurat object and an updated matrix for each step. The user can easily manipulate the Seurat object by loading it into R. Since cluster annotation has not been integrated into the pipeline, users can manually annotate the clusters using marker genes or cell type reference signatures.

## 6 Availability

`scAN1.0` is freely available at: https://gitbio.ens-lyon.fr/LBMC/sbdm/scan10. Please, take caution to clone using HTTP instead of SSH.

## 7 Acknowledgements

We thank the Institut Convergence Plascan (Grant Number ANR-17-CONV-0002) for their support.

Mirela Diana Ilie has been supported by the Fondation ARC pour la recherche sur le cancer (ARCMD-DOC12020020001361).

We thank (i) Emmanuel Jouanneau and Alexandre Vasiljevic from the Neurosurgery and Pathology Departments, Reference Center for Rare Pituitary Diseases HYPO, “Groupement Hospitalier Est” Hospices Civils de Lyon, Bron, France for providing diagnosed Gonadotroph tumor material; (ii) Hector Hernandez-Vargas for his advices and support with setting the single cell dissociation and subsequent bioinformatic analysis; and (iii) Laurent Modolo from the LBMC for his help using the Nextflow syntax.

We gratefully acknowledge support from the PSMN (Pôle Scientifique de Modélisation Numérique) of the ENS de Lyon for the computing resources

We finally thank the BioSyL Federation (http://www.biosyl.org), the LabEx Ecofect (ANR-11-LABX-0048) and the LabEx Milyon of the University of Lyon for inspiring scientific events.

## 8 Author contributions

Maxime Lepetit wrote the `scAN1.0` script, performed the analysis, generated the figures and wrote the paper. Mirela Diana Ilie and Gerald Raverot participated in generating and analyzing the Gonadotroph tumour dataset. Marie Chanal participated in generating the Gonadotroph tumour dataset. Christophe Arpin participated in analyzing the CD8 T cells dataset and in writting the paper. Philippe Bertolino, Franck Picard and Olivier Gandrillon helped designing the study and analyzing the results, wrote the paper, and secured the funding.

## References

[1] Y. Hao, S. Hao, E. Andersen-Nissen, 3rd Mauck W. M., S. Zheng, A. Butler, M. J. Lee, A. J. Wilk, C. Darby, M. Zager, P. Hoffman, M. Stoeckius, E. Papalexi, E. P. Mimitou, J. Jain, A. Srivastava, T. Stuart, L. M. Fleming, B. Yeung, A. J. Rogers, J. M. McElrath, C. A. Blish, R. Gottardo, P. Smibert, and R. Satija. “Integrated analysis of multimodal single-cell data”. In: Cell 184.13 (2021), 3573–3587 e29.

[2] P. Di Tommaso, M. Chatzou, E. W. Floden, P. P. Barja, E. Palumbo, and C Notredame. “Nextflow enables reproducible computational workflows”. In: Nature biotechnology 35.4 (2017), pp. 316–319.

[3] T. Chari and L. Pachter. “The Specious Art of Single-Cell Genomics”. In: bioRxiv (2022), p. 2021.08.25.457696.

[4] N. S. Kurd, Z. He, T. L. Louis, J. J. Milner, K. D. Omilusik, W. Jin, M. S. Tsai, C. E. Widjaja, J. N. Kanbar, J. G. Olvera, T. Tysl, L. K. Quezada, B. S. Boland, W. J. Huang, C. Murre, A. W. Goldrath, G. W. Yeo, and J. T. Chang. “Early precursors and molecular determinants of tissue-resident memory CD8(+) T lymphocytes revealed by single-cell RNA sequencing”. In: Sci Immunol 5.47 (2020).

[5] R. S. Brüning, L. Tombor, M. H. Schulz, S. Dimmeler, and D. John. “Comparative analysis of common alignment tools for single-cell RNA sequencing”. In: Gigascience 11 (2022).

[6] R. Satija, J. Farrell, and D. Gennert. “Spatial reconstruction of singlecell gene expression data.” In: Nat Biotechnol 33.5 (2015), pp. 495–502.

[7] C. Hafemeister and R. Satija. “Normalization and variance stabilization of single-cell RNA-seq data using regularized negative binomial regression.” In: Genome Biol 20.1 (2019), p. 296.

[8] A.T. L. Lun, K. Bach, and J.C. Marioni. “Pooling across cells to normalize single-cell RNA sequencing data with many zero counts.” In: Genome Biol 17.1 (2019), p. 75.

[9] P. Melsted, A. S. Booeshaghi, F. Gao, E. Beltrame, L. Lu, K. E. Hjorleifsson, J. Gehring, and L. Pachter. “Modular and efficient preprocessing of single-cell RNA-seq”. In: Nat. Biotechnol. 39 (2021), pp. 813–818.

[10] D. A. Jackson. “Stopping Rules in Principal Components Analysis: A Comparison of Heuristical and Statistical Approaches”. In: Ecology 74 (1993), pp. 2204–2214.

[11] S. Chen, Y. Zhou, Y. Chen, and J. Gu. “fastp: an ultra-fast all-in-one FASTQ preprocessor”. In: Bioinformatics 34.17 (2018), pp. i884–i890.

[12] G. X. Zheng, J. M. Terry, P. Belgrader, P. Ryvkin, Z. W. Bent, R. Wilson, S. B. Ziraldo, T. D. Wheeler, G. P. McDermott, J. Zhu, M. T. Gregory, J. Shuga, L. Montesclaros, J. G. Underwood, D. A. Masquelier, S. Y. Nishimura, M. Schnall-Levin, P. W. Wyatt, C. M. Hindson, R. Bharadwaj, A. Wong, K. D. Ness, L. W. Beppu, H. J. Deeg, C. Mc-Farland, K. R. Loeb, W. J. Valente, N. G. Ericson, E. A. Stevens, J. P. Radich, T. S. Mikkelsen, B. J. Hindson, and J. H. Bielas. “Massively parallel digital transcriptional profiling of single cells”. In: Nat Commun 8 (2017), p. 14049.

[13] T. Ilicic, J.K. Kim, A.A. Kolodziejczyk, F.O. Bagger, D.J. McCarthy, J.C. Marioni, and Teichmann S.A. “Classification of low quality cells from single-cell RNA-seq data.” In: Genome Biology 17.1 (2016), p. 29.

[14] D. J. McCarthy, K. R. Campbell, A. T. L. Lun, and Q. F. Wills. “Scater: pre-processing, quality control, normalization and visualization of single-cell RNA-seq data in R”. In: Bioinformatics 33.8 (Jan. 2017), pp. 1179–1186. eprint: https://academic.oup.com/bioinformatics/article-pdf/33/8/1179/25150420/btw777.pdf.

[15] C.S. McGinnis, L.M. Murrow, and Z.J. Gartne. “DoubletFinder: doublet detection in single-cell RNA sequencing data using artificial nearest neighbors.” In: Cell systems 8.4 (2019), 329–337. e4.

[16] N.M. Xi and J.J. Li. “Benchmarking Computational Doublet-Detection Methods for Single-Cell RNA Sequencing Data”. In: Cell Systems 12.2 (2021), 176–194.e6.

[17] J. Breda, M. Zavolan, and E. van Nimwegen. “Bayesian inference of gene expression states from single-cell RNA-seq data”. In: Nat Biotechnol (2021).

[18] R. Zhang, G. S. Atwal, and W. K. Lim. “Noise regularization removes correlation artifacts in single-cell RNA-seq data preprocessing”. In: Patterns (N Y) 2.3 (2021), p. 100211.

[19] V. D. Blondel, J.-L. Guillaume, R Lambiotte, and E. Lefebvre. “Fast unfolding of communities in large networks”. In: J. Stat. Mech. P10008 (2008), p. 12.

[20] Peter J. Rousseeuw. “Silhouettes: A graphical aid to the interpretation and validation of cluster analysis”. In: Journal of Computational and Applied Mathematics 20 (1987), pp. 53–65.

[21] L. Van der Maaten and G. Hinton. “Visualizing data using t-SNE”. In: Journal of Machine Learning Research 9 (2008), pp. 2579–2605.

[22] L. McInnes, J. Healy, N. Saul, and L. Großberger. “UMAP: uniform manifold approximation and projection”. In: J. Open Source Softw 3 (2018), p. 861.

